# *In vivo* contactless brain stimulation via non-invasive and targeted delivery of magnetoelectric nanoparticles

**DOI:** 10.1101/2020.05.29.123075

**Authors:** Tyler Nguyen, Jianhua Gao, Ping Wang, Abhignyan Nagesetti, Peter Andrews, Sehban Masood, Zoe Vriesmann, Ping Liang, Sakhrat Khizroev, Xiaoming Jin

## Abstract

Non-invasive brain stimulation is valuable for studying neural circuits and treating various neurological disorders in humans. However, the current technologies usually have low spatial and temporal precision and poor brain penetration, which greatly limit their application. A new class of nanoparticles known as magneto-electric nanoparticles (MENs) can be navigated to a targeted brain region with a magnetic field and is highly efficient in converting an externally applied magnetic wave into local electric fields for neuronal activity modulation. Here we developed a new method to fabricate MENs of CoFe_2_O_4_-BaTiO_3_ core-shell structure that had excellent magneto-electrical coupling properties. Using calcium imaging of organotypic and acute cortical slices from GCaMP6s transgenic mice, we demonstrated their efficacy in reliably evoking neuronal responses with a short latency period. For *in vivo* non-invasive delivery of MENs to brain, fluorescently labeled MENs were intravenously injected and guided to pass through the blood brain barrier to a targeted brain region by applying a magnetic field gradient. A magnetic field (∼450 Oe at 10 Hz) applied to mouse brain was able to reliably evoke cortical activities, as revealed by *in vivo* two-photon and mesoscopic imaging of calcium signals at both cellular and global network levels. The effect was further confirmed by the increased number of c-Fos expressing cells after stimulation. Neither brain delivery of MENs nor the subsequent magnetic stimulation caused any significant increases in the numbers of GFAP and IBA1 positive astrocytes and microglia in the brain. This study demonstrates the feasibility of using MENs as a novel efficient and non-invasive technique of contactless deep brain stimulation that may have great potential for translation.

## Introduction

In the last two decades, noninvasive brain stimulation techniques have made important contributions to the understanding of brain neurophysiology and treatment of neurological diseases. The two most common forms of noninvasive brain stimulation used clinically are Transcranial Magnetic Stimulation (TMS) and transcranial Direct Current Stimulation (tDCS)^1, 2, 3^. However, they have relatively low spatial and temporal precisions, which greatly diminish their efficacy and application. For example, regular TMS has a spatial resolution of 3-5 cm and depth of penetration of ∼1-1.5 cm, which allows for stimulating a cortical gyrus^4, 5^. tDCS also has very low spatial resolution and limited effects on deep brain structures^4, 6, 7^. More recent approaches such as optogenetics^8, 9^, magneto-thermal brain stimulation^10, 11^, and ultrasound-based transcranial stimulations^1212, 13,14, 14^, have improved in ways that were limited in tDCS and TMS (i.e. cellular specificity and spatial and temporal resolutions) but present drawbacks including the requirement of genetic modifications (e. g. optogenetics), invasive brain delivery of particles (e.g. magneto-thermal nanoparticles), and brain tissue damage due to high release of thermal energy (e.g. in focused ultrasound brain stimulation).

The concept of using magneto-electric nanoparticles (MENs) for brain stimulation was first proposed in a theoretical paper by Yue et al in 2012^15^. The same research group led by Sakhrat Khizroev reported the first in vivo MEN-mediated modulation of brain EEG activity by external magnetic field via intravenous injection of MENs in 2015 ^16^. They used coreshell CoFe_2_O_4_-BaTiO_3_ MENs due to their relatively strong magneto-electric (ME) effect owing to the strain-induced coupling between the magnetostrictive cobalt ferrite cores and piezoelectric barium titanate shells. The nanoparticles can be injected into a vein, forced to cross blood-brain barrier (BBB) and consequently localized to a target region by applying a magnetic field gradient. When exposed to a relatively low magnetic field (100-500 Oe), the MEN’s cobalt ferrite (CoFe_2_O_4_) core experiences non-zero strain due to the magnetostrictive effect. Owing to the ME coupling, this strain propagates through the lattice matched interface to the adjacent barium titanate (BaTiO_3_) shell, which in turn induces a local electric field (on the order of 1000 V/m) due to the piezoelectric effect ^16^. In this way, magnetic brain stimulation using MENs (MENs-MS) can be achieved with a magnetic field intensity that is much lower than what is used for typical TMS (30,000-50,000 Oe)^17^. The unique properties of MENs, due to their small size (∼30 nm) and the ME effect not displayed by any other nanoparticles known to date, may provide significant improvements over currently used techniques in in efficacy, precision, and tissue penetration for noninvasive brain stimulation. However, to date, no direct effect of MENs on stimulating neurons in a focal brain region and/or a large neural network has been demonstrated.

In the current study, we developed a non-invasive technique using MENs to wirelessly stimulate cortical neuronal activity. Using *in vitro* and *in vivo* fluorescent and two-photon imaging techniques, we showed that MENs can be drawn to across the BBB and localized to a target cortical region without any apparent signs of neuroinflammation. By activating the MENs with a weak external magnetic field at a specific frequency, we were able to induce cortical activities in individual neurons and in a large neural network *in vivo*.

## Procedures

### 1. MENs synthesis, ultrastructural imaging, and functionalization

MENs were synthesized using the hydrothermal method as described in Guduru et al. 2014 ^18^. Samples for transmission electron microscope (TEM) imaging were prepared by dispersing 1 mg MENs in 2 mL ethanol and sonicating the mixture for 20 minutes. One drop of the MENs solution was deposited with a pipette on a copper TEM grid substrate. Slowing evaporation at low temperature provided a uniform nanoparticle distribution on the substrate. To eliminate aggregation and potential toxicity, the surfaces of the MENs were functionalized by adding a 2-nm thick coating of glycerol mono-oleate (GMO) as follows: the MENs were washed twice with phosphate-buffered saline (PBS, pH 7.4) and sonicated for 2 minutes between washes for uniform dispersion in solution. MENs were then suspended in a GMO solution (8 µL GMO/1 g MENs) and mixed thoroughly on a rotator for 12 hours. Excess GMO were then removed by extraction with 1 ml of 70% ethanol: PBS and two subsequent washes with PBS. Finally, the MENs-GMO mixture were suspended in saline and stored at −20 °C for short-term storage of 3-5 days or freeze dried in liquid nitrogen for long term storage for more than 1 week. To prepare fluorescent MENs (fMENs), Texas Red NHS ester was first reacted with oleylamine (2:1 molar ratio) for 18-24 hours on a rotator in a dark environment. Then, hexane was added in drops incrementally to help dissolved the oleylamine-Texas Red mixture. After the solution of oleylamine-Texas Red NHS ester was combined with the GMO-MENs mixture (1:3 volume ratio of GMO-MENs: oleyamine-TexasRed), it was rotated for additional 12 hours to allow binding. Following removal of excess GMO through multiple washes in PBS, the final product was store at −20 °C (short-term storage).

### 2. *In vitro* imaging of cortical neurons in organotypic cultures from GCaMP6s transgenic mice

All experiments were approved by the Institutional Animal Care and Use Committee (IACUC) of the Indiana University School of Medicine, which are in accordance with National Institutes of Health guidelines for the care and use of laboratory animals. *<u>Preparation of cortical slice cultures:</u>* Cortical slice cultures were prepared from GCaMP6s transgenic mice that were 3-5 days old (n=5). The animals were anesthetized by rapidly cooling them in ice. After quick decapitation, the brain was removed and immediately immersed in an ice-cold cutting artificial cerebrospinal fluid (aCSF) solution (2 mM KCl, 1 mM MgCl_2_, 2 mM MgSO_4_, 1.25 mM NaH_2_PO_4_, 26 mM NaHCO_3_, 1 mM CaCl_2_, 10 mM D-Glucose, 206 mM Sucrose). The cerebellum was then removed, and the brain was glued onto a cutting stage. A vibratome (VT1200, Leica Biosystems, Buffalo Grove, IL.) was used to section the brain into coronal slices of 350 µm thick at a cutting velocity of 0.75 mm/minutes. The slices were then collected and placed in a cold oxygenated aCSF solution (126 mM NaCl, 2.5 mM KCl, 2 mM MgSO_4_, 1.25 mM NaH_2_PO_4_, 26 mM NaHCO_3_, 2 mM CaCl_2_, 10 mM D-Glucose). Intact slices were then selected and arranged carefully on Millipore Millicell culturing inserts (Millipore Sigma, St. Louis, MO). The slices were cultivated at ∼37 °C in 95% O_2_/5% CO_2_ in a culturing medium consisting of 46% Basal Medium Eagle (BME), 25% Earle’s Salt Solution, 25% fetal bovine solution (FBS), 3% of 22% Glucose solution, and 1% L-Glutamine-Penicillin-Streptomycin solution. 75% of the medium was changed with new medium every three days.

#### Magnetic field application

Prior to imaging, a cultured brain slice was removed from the culturing insert by cutting the membrane surrounding the slice and placed in a custom-made recording chamber filled with 37 °C aCSF. The recording chamber was a 2-cm diameter well with a glass bottom that was filled with aCSF containing 5 µL of MENs (at a final concentration of 5 mg/mL). A 5000 Oe conical magnet was used to apply a strongly localized magnetic field to the bottom of the chamber for approximately 15 minutes to draw the MENs to a focal area of the slice. For magnetic stimulation, two electromagnets were put on opposite sides of the recording chamber in a Helmhotz pair arrangement according wherein the diameter of each coils was comparable to the separation between the coils. This arrangement was chosen to significantly reduce or eliminate a magnetic force to ensure the nanoparticles did not physically move during the stimulation process. A sinusoidal magnetic field of approximately 450 Oe was applied in a pulsed mode with 50 ms pulse width at 10 Hz to stimulate the MENs during the imaging process.

#### Two-photon imaging

Two-photon imaging of cortical slices was performed using a Prairie Technologies Ultima 4423 two-photon system (Bruker Inc., Middleton, WI) equipped with a MaiTai Ti: Sapphire laser (Newport, Mountain View, CA) tuned to 900 nm. Band-pass filtered fluorescence (560-600 nm) was collected by photomultiplier tubes of the system. The average laser power on the sample was ∼20-30 mW. All images were acquired at a resolution of 512 x 512 pixels using a 20x water-immersion objective (Nikon Instruments Inc., Melville, NY). Images were capture at a rate of 4-5 fps. Typically, each recording session consisted of 30 seconds of baseline recording, 40s with magnetic field application, and 30 seconds of post-magnetic application period.

#### *Ex vivo* calcium imaging of acute GCaMP6s cortical slices

*<u>Cortical slice preparation</u>*:Thy1-GCaMP6s transgenic mice, in which layers II/III and V pyramidal neurons express GCaMP6s calcium sensor protein, were purchased from the Jackson’s Laboratory and bred locally. Adult mice (n=4) were anesthetized with 78 mg/kg ketamine and 22 mg/kg xylazine (i. p.) and decapitated. The brain was removed and immediately placed in an ice-cold oxygenated sucrose artificial cerebral spinal fluid (s-ACSF) solution (206 mM sucrose, 2 mM KCl, 1 mM MgCl2, 2 mM MgSO4, 1.25 mM NaH2PO4, 26 mM NaHCO3, 10 mM D-glucose, 1 mM CaCl2). After 1-2 minutes, the brain was situated on a cutting stage with the cortex facing the approaching blade. Slices of 350 µm thickness were cut with a Vibratome (Leica VT1200S; Leica, Nusslock, Germany) while the brain was submerged in the cutting solution. The slices were then incubated at 37 °C for 1 hour in a chamber filled with oxygenated ACSF (124 mM NaCl, 3 mM KCl, 2 mM MgSO4, 1.25 mM NaH2PO4, 26 mM NaHCO3, 10 mM D-glucose, 1 mM CaCl2). *<u>Calcium imaging:</u>* A recording chamber was made by attaching a temperature control heated well (Thermal Well Temperature Controller TC-100, BioScience Tools, San Diego, CA) onto a coverslip. Liquid inflow and vacuum outflow tubing were installed to generate ACSF current through the chamber. A cortical slice was placed in the chamber and secured with a metal ring. Calcium activities of the slices was imaged with a system consisting of a Leica DM6000 FS upright microscope with a fluorescence light source directed through a 10x water immersion objective (Leica, Nusslock, Germany). Images were capture with an iXON EMCCD DU-88U camera system (Andor USA, Concord, MA) at ∼45-50 fps. Cortical slices were imaged at ∼35 °C in aCSF with 20 µM bicuculline.

#### MENs loading and stimulation parameters

After the liquid flow of the recording chamber was turned off briefly, 20 µL of MENs (concentration of 5 mg/mL in aCSF) was added to the chamber. A 5000 Oe conical magnet was applied under the glass for ∼5-8 minutes to draw MENs to a small area of the slice. Liquid flow was turned back on prior to stimulation and recording. For stimulation, the recording chamber was placed between a pair of electromagnets. A unipolar magnetic field of 750-875 Oe with 200 ms pulse-width was applied during the imaging process.

### 3. *In vivo* two-photon imaging of fMENs in cerebral blood stream

Thy1-GCaMP6s mice (n=5) were anesthetized with Ketamine/Xylazine mixture (87.3 mg/kg / 13.7 mg/kg) and the scalp was removed. A cranial window (3 mm in diameter) was made in an area 1.5 mm lateral from the midline and 1 mm posterior from the bregma. Two pieces of cover glass (a 3-mm diameter glass adhered to a 5-mm diameter glass) were glued together with cyanoacrylate glue with the 3-mm glass facing the brain surface. After the glass was installed on the cranial window, an L-shaped titanium head-plate was glued on the posterior region of the head for head fixation during imaging. After the animals were allowed to recover for 7 to 10 days, a single injection of 200 µl of fMENs at 200 µg/ml was made through the retro-orbital route. Two-photon images of cerebral blood vessels were taken at baseline, and 0, 5, 10, 20, and 30 minutes after fMENs injection.

#### *In vivo* two-photon imaging after fMENs delivery

In GCaMP6s mice with cranial windows prepared as described above, an injection of 200 µL fMENs at 5 mg/ml was made through the retro-orbital route. After two sets of stacked cylindrical magnets of 6000 Oe field strength were separately placed on top of the cranial window and beneath the mouth of the mouse for 10 minutes, a conical magnet with the field of 5000 Oe at the tip was applied to the cranial window to further localize the MENs. Images were taken before and after fMENs injection, after fMENs injection with magnet application, and 24 hours after MEN delivery.

### 4. *In vivo* two-photon calcium imaging of cortical neurons in GCaMP6s transgenic mice

Mice with cranial windows prepared as described above were used in this experiment. After the animals were sedated via 4% isofluorane and maintained with intraperitoneal injection of chloprothixene (0.04 mg/ml), they were injected retro-orbitally with 200 µl MENs at 200 µg/ml. Following the delivery of the MENs to cortex using the same technique described above, the animal’s head was stabilized by fixing the L-shaped titanium metal plate to a custom-made base and the body temperature was maintained. Magnetic stimulation was made by placing two electromagnets (∼500 ms pulse-width at ∼300-450 Oe) closely on both side of the head. Again, the Helmholtz pair arrangement of the electromagnet was chosen to minimize the pulling magnetic force on the nanoparticles during the stimulation events. Two-photon images of calcium transients of layer II/III neurons were taken at baseline and during and after magnetic stimulation. In each imaging field, two optical planes in layer II/III were imaged for 2 minutes at 4-5 frames per second (fps). We tested a combination of different stimulation frequencies (5, 10, 20, 50, and 100 Hz).

### 5. Mesoscopic calcium imaging of cortical activity *in vivo*

Thy1-GCaMP6s transgenic mice were anesthetized with Ketamine/Xylazine mixture (87.3 mg/kg / 13.7 mg/kg) and the scalp was removed. After a large cranial window (approximately 10 x 8 mm) was made by removing the skull, a piece of curved cover glass was installed and tightly sealed with super glue and dental cement. The animals were allowed to recover for 7 to 10 days prior to imaging. A mesoscopic imaging set-up was built similar to previously described method^19, 20^. Images were captured with a two-lens system composed of a top zoom Nikkor 70-300 mm lens of aperture f3.5-5.6 set at 70mm at f3.5 coupled with an inverted lens adapter to a bottom Nikkor 50 mm prime lens at f1.4. Videos were captured with an iXON EMCCD DU-88U camera system (Andor USA, Concord, MA) controlled by MetaMorph software. Delivery of MENs and magnetic stimulation paradigm were done exactly as described in two-photon imaging.

### 6. Data analysis of Ca imaging

Two-photon imaging data were analyzed post hoc using ImageJ software (NIH) with a Time Series Analyzer plugin (Balaji J. UCLA). All active cells (cells that produce calcium fluorescence flashes throughout all frames) within each frame were analyzed. To analyze individual cells, three separate background points were chosen by drawing circles with the same diameter as the respective cell body in the surrounding area. The background areas selected did not include dendrites and other visible neuronal structures. Data obtained included signals of cell body and three background fluorescent profiles. For mesoscopic brain imaging, a circular area of interest (∼1 mm in diameter) was selected, with an adjacent blood vessel selected as background to compensate for changes in general brightness. The fluorescence calcium signal of a cell ΔF/F was calculated by equation (1) excerpt from Chen et al. 2013 ^21^ that was developed by Kerlin et al, 2010^22^:

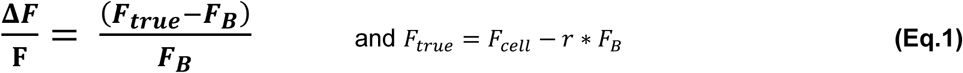

where F_B_ is the average baseline fluorescence over three different regions surrounding the cell of interest and r is the contamination ratio standard constant 0.7. The data were then used to construct a calcium peak profiles using OriginPro 9.1 (OriginLab, Northampton, MA). Analyses of peak amplitude, frequency, and durations were done with Peak analysis toolbox of OriginPro 9.1.

### 7. C-fos staining of cortical neurons

C57BL6J mice (n = 4 animals/group) were assigned into three groups: control (saline injection + MS (5 x 2 minutes MS at 10 Hz)), MENs (MENs delivery without magnetic stimulation), and MENs-MS (MENs delivery and MS (5 x 2 minutes MS at 10 Hz)). MENs delivery and magnetic stimulation (MS) were done similarly as described above.

Brain tissues of these mice were stained for c-Fos protein, an indicator of action potential firing^23^. Four hours after the treatment, the mice were deeply anesthetized and perfused transcardially with PBS buffer, followed by 4% formaldehyde PFA at room temperature. The brains were then dissected and placed in 30% sucrose for 48-72 hours at 4°C. The tissue was then frozen and sectioned with a cryostat. Immunofluorescence was performed on free-floating sections by incubating overnight with a primary antibody for c-Fos (rabbit; 1:800, Sigma Aldrich G3893) and followed by incubation with fluorescent goat anti-rabbit secondary antibody (1:200; Invitrogen, Carlsbad, CA). For nuclear staining, 4’, 6-diamidino-2-phenylindole (DAPI; 1: 10,000) was added to the solution for a final 5 min. Finally, slices were imaged using a Neurolucida imaging system.

### 8. Immunofluorescence analysis of astrocytes and microglia

To determine potential glial reactivity and neuroinflammation induced by the MENs, we used an antibody against ionized calcium binding adaptor protein (IBA1) to label microglia, and an antibody against Glial Fibrillary Acidic Protein (GFAP) to label astrocytes^24, 25^. C57BL6J mice were divided into three groups: control (saline +MS), MENs (MENs without MS), and MENs-MS (MENs delivery with MS). The MENs delivery and MS paradigm were identical to the c-Fos experiment described above. Within the MENs-MS group, subgroups of mice were euthanized at 4 hours, 24 hours, 3 days, or 1 week after MS. Mice were perfused and brain slices were stained as described above. Anti-GFAP (mouse; 1:800, Sigma Aldrich G3893) and anti-IBA1 (goat; 1:200, ABCam ab5076) antibodies were followed by secondary antibodies of goat anti-mouse Cy5 (1:500, Jackson Immuno) and donkey anti-goat 488nm (1:1000, Fischer Scientific).

### 9. Statistics

Mean values and final plots were developed in Microsoft Excel, Jmp Analysis 11 (SAS Institute Inc. 2013. Cary, NC), and GraphPad Prism 6 (GraphPad Software, La Jolla, California). All statistical analyses were done with Jmp Analysis 11 and GraphPad Prism 6. ANOVA analyses were used for the following comparisons: calcium amplitudes of the same group across different time points (Repeated measures Anova), amplitudes and frequencies of calcium imaging data (one-way), calcium amplitudes and frequencies across different tested magnetic frequencies and different cortical region in mesoscopic imaging (one-way), and compare of positive stained cells between groups of c-Fos, IBA1, and GFAP immunohistochemistry experiment (one-way). For comparisons that yielded statistical significances, Tukey’s HSD post-hoc analyses were applied for further comparisons between specific groups.

## Results

### 1. MENs synthesis and characterization

We developed MENs with a true coreshell nanostructure, in which the magnetic core was intrinsically coupled to the piezoelectric shell via perfect lattice matching (Figure 1). The size of the MENs was on the order of 30 nm. With this new synthesis, we were able to develop MENs with a high ME coefficient of >5 V/cm/Oe in the frequency range, 0 to 200 Hz, under study (data not shown).

**Figure 1.**
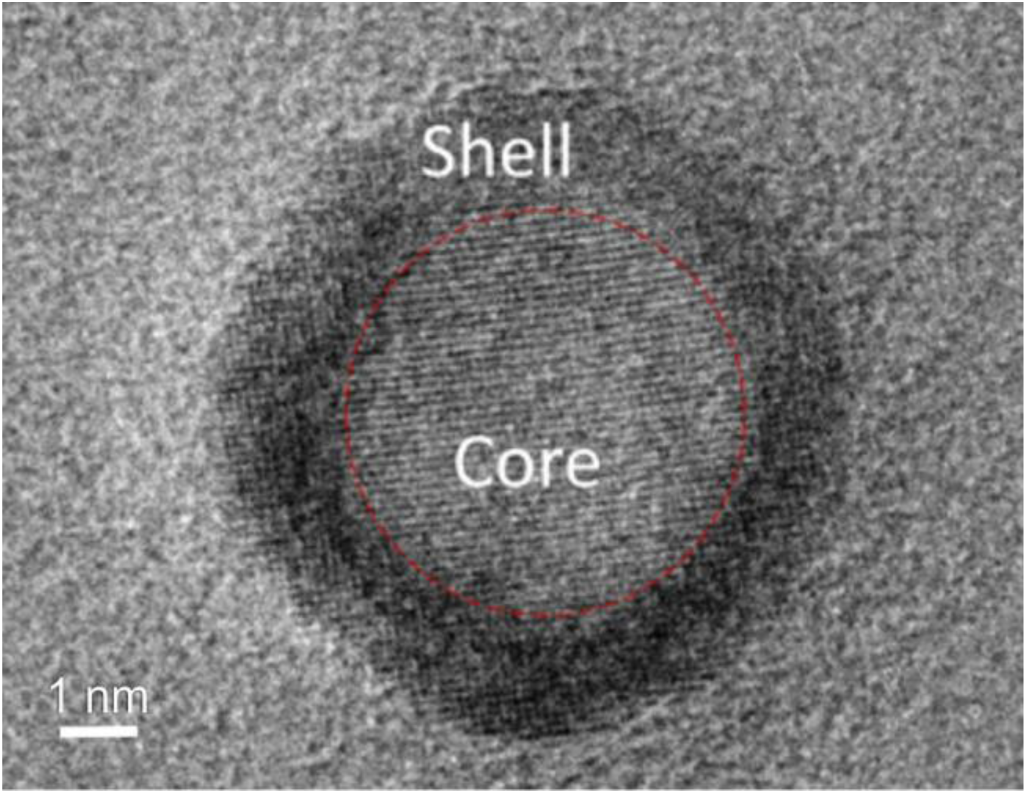
Transmission electron microscopy (TEM) image of a magnetoelectric nanoparticle (MEN). A TEM image demonstrates an almost-perfect lattice matched surface interface between the magnetostrictive core and the piezoelectric shell (shown with broken line) of the coreshell nanostructure.

### 2. MENs-MS induced neuronal activity in cultured and acute cortical slices

We first assessed the effects of MENs-MS on neuronal activity in organotypic cortical slices prepared from GCaMP6 transgenic mice. After the slices were cultured for 5 days old *in vitro*, they were loaded with MENs and stimulated with a magnetic field (10 Hz sinusoidal wave at 450 Oe for 10s). The peak amplitude of calcium transients from individual neurons increased nearly 200% (Figure 2A and B, F/F_o_ 0.809 ± 0.12 for baseline vs 3.586 ± 0.78 for magnet-on, p < 0.005, Repeated-measures ANOVA, Tukey’s HSD). In neurons that were initially active, we saw a significant increase in calcium signals, whereas in neurons that were not active, the magnetic stimulation induced calcium activity (Figure 2A and B). We also found a significant increase in spike frequency during the period when the MS was turned on (Figure 2C and D, 1.43 ± 0.13, 2.6 ± 0.21, and 1.46 ± 0.17 spikes/cell for baseline, MS-on, and MS-off periods, respectively, p < 0.01 for comparison between baseline and MS-on, Repeated ANOVA, Tukey’s HSD).

**Figure 2.**
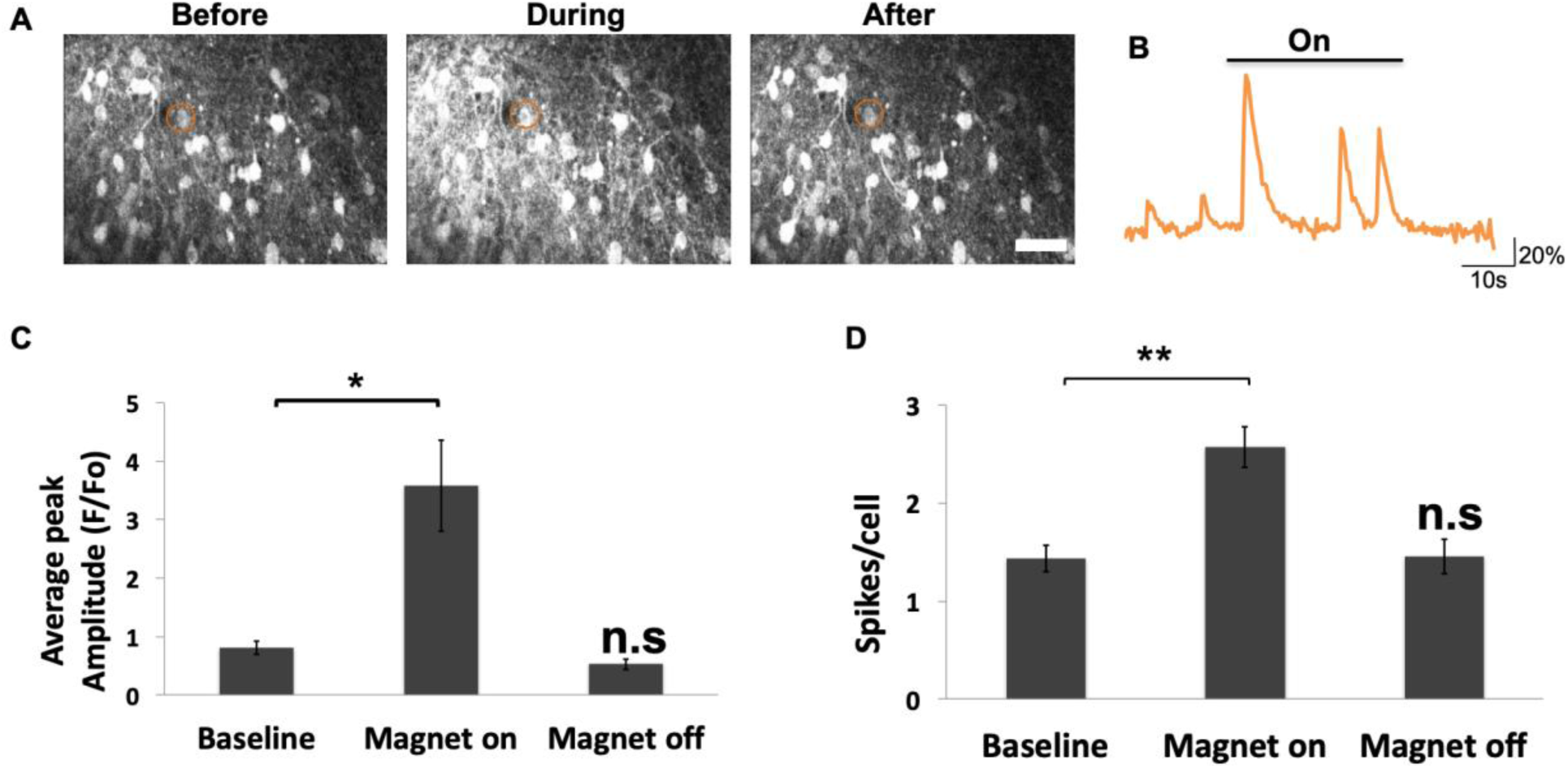
Magnetic stimulation of MENs-loaded cortical slice cultures induced neuronal activity. Cultured cortical slices that expressed GCaMP6 in pyramidal neurons were loaded with MENS by adding 20 µl of 5 mg/mL MENs solution, followed by applying a conical permanent Neodymium (Nd) magnet for ∼15 minutes. Magnetic stimulation was made with a pair of electromagnets at 10 Hz. A. Representative two-photon images of calcium signals in periods before, during, and after magnetic stimulation. B. A sample trace of calcium transients measured from a neuron labeled with red circles in (A). **C, D**. Increases in average calcium spike amplitude (C) and number of spikes per cell (D) during the stimulation period. Calcium activity returned to baseline level after the magnet was off. *<u>Scale bar</u>*: 50 µm. *: p<0.05, **: p<0.01, Repeated measured ANOVA, Tukey’s HSD. n = 9 slices.

In order to improve the temporal precision of the stimulation, we used a single pulse (unipolar square pulse at 750-875 Oe, 200 ms duration) of magnetic field and tested the effect in acutely prepared cortical slices from GCaMP6 transgenic mice (Figure 3A). The mean amplitude of calcium spikes at baseline was 0.127 ± 0.035, which became 0.541 ± 0.056 during magnetic stimulation, and returned to 0.121 ± 0.32 after the magnet was turned off (p < 0.005, Repeated ANOVA, Tukey’s HSD; Figure 3B). Most of the evoked calcium responses had a shorter latency period (Figure 3C), with over 80% of the responses occurring in less than 1 second and 32% of the responses in less than 150 milliseconds after the magnetic pulse was turned on.

**Figure 3.**
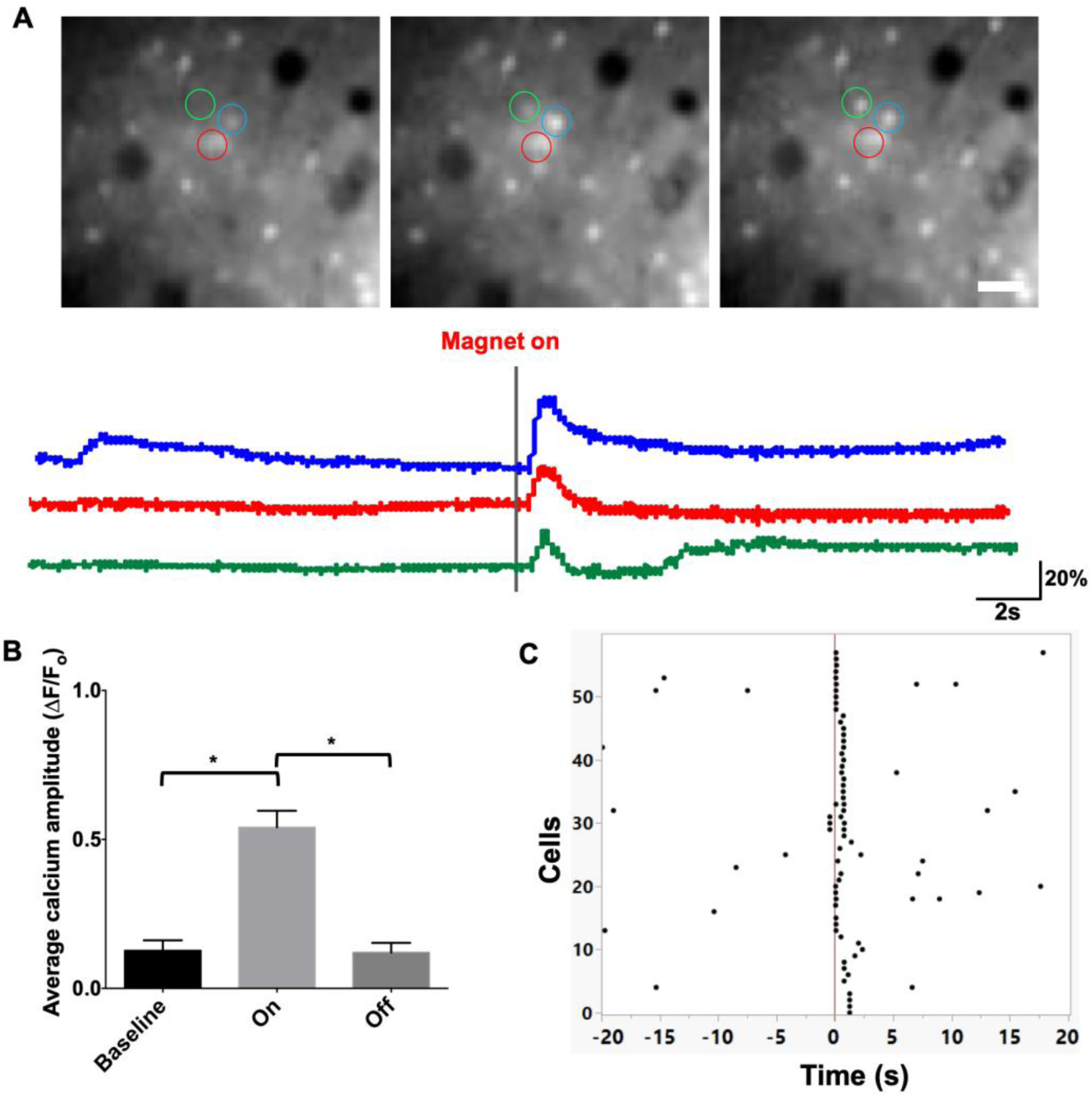
Single unipolar magnetic pulses induced neuronal activity in MENs-loaded cortical slices. Cortical slices prepared from GCaMP6 transgenic mice were loaded with MENs similar to Fig. 2 and stimulated with single pulses of magnetic field (∼750-875 Oe at 200 ms pulse-width). **A.** Sample images of neurons in a slice and traces of calcium transients measured from the circled neurons. **B**. Average peak amplitudes of calcium spikes measured at baseline and after the magnets were turned on and off. **C.** Temporal response profile of all responding cells. *<u>Scale bar:</u>* 50 µm, *p<0.005, Repeated-measured ANOVA, Tukey’s HSD. n = 13 slices (total 4 animals).

### 3. Intravenous injection of MENs followed by magnetic delivery to brain

We first use repeated *in vivo* two-photon imaging of cerebral blood circulation to determine how long intravenously injected MENs would be detectable in blood stream. Immediately after injecting fMENs into the tail vein of a mouse, we found many fluorescent particles in blood stream of cortical vessels. These particles decreased rapidly over a short period of time (Figure 4A-B): with the fluorescence intensity in blood stream dropping to about a half in ∼15 minutes and to almost zero in ∼30 minutes after injection (Figure 4B). This drop-in particle number suggested that the optimal time window for applying a magnet for MENs delivery is within ∼10 minutes after intravenous injection of MENs.

**Figure 4.**
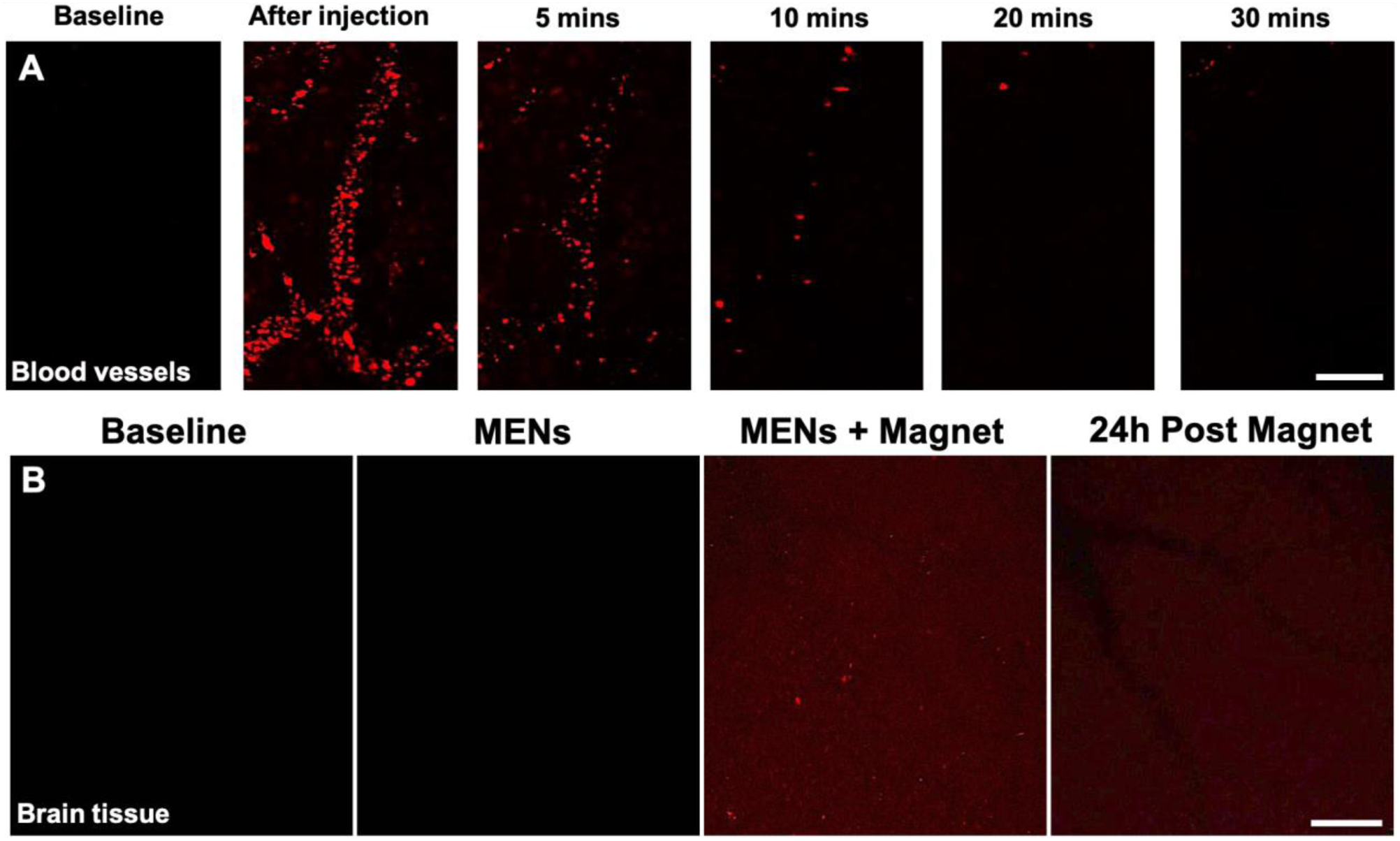
*In vivo* delivery of fMENs into mouse cortex. **A.** Vascular MENs level reduced significantly after 10 minutes post injection. *In vivo* two-photon images of cortical blood vessels were taken at baseline (B.L.) and at 5, 10, 20, and 30 minutes after intravenous injection of 10 µg fluorescence labeled MENs (fMENs). No magnetic field was applied. *<u>Scale bar:</u>* 100 µm. **B**. *In vivo* two-photon images of layer 2/3 sensorimotor cortex were taken at baseline (C1) and after (C2) intravenous injection of 10 µg fMENs, after applying ∼4000-5000 Oe magnetic field for 45 minutes (C3), and on the second day after injection (C4) indicates successful delivery of MENs to cortical tissue after magnet application, which MENs remained up to at least 24 hours after delivery. *<u>Scale bar:</u>* 100 µm.

We used fMENs to assess the effectiveness of applying permanent magnet for delivering MENs into brain parenchyma. Following intravenous injection of fMENs and application of a conical magnet on mouse head for 20 minutes (Figure 4C-D), *in vivo* two-photon imaging showed that fluorescence signals were visible through cranial windows in mice with fMENs delivery, but not in mice with MENs injection only. The result suggests that the fMENs were drawn by a magnetic field to cross BBB and entered brain parenchyma *in vivo* (Figure 4C-D). When the same mice were imaged again 24 hours later, significant signals of MENs were still evident in the same region (Figure 4D), suggesting successful delivery of MENs to cortical tissue.

### 4. MENs-MS induced cortical neuronal activity and network activity *in vivo*

We used *in vivo* two-photon imaging in GCaMP6 transgenic mice to assess whether MENs-MS would evoke cortical neuronal activity (Figures 5A-D). Following delivery of MENs and MS with a 10 Hz magnetic wave, we found that there was a dramatic increase in the mean amplitude of calcium spikes (Figure 5C. F/F_o_ of 1.50 ± 0.09, 2.37 ± 0.18, and 1.36 for baseline, MS-on, and MS-off, respectively, p<0.05, Repeated-measures ANOVA, Tukey’s HSD), and spike frequency (3.08 ± 0.27 spikes/cell vs. 1.96 ± 0.24 for baseline with p=0059 and 1.54 ± 0.25 for magnet off with p<0.05, repeated ANOVA, Tukey’s HSD; Figure 5D) in cortical layer II/III neurons. The efficacies of magnetic waves at 5, 10, 20, 50, and 100 Hz on activating cortical neurons were also tested *in vivo*. We found that, for every magnetic frequency applied during the stimulation period, a significant increase in somatic calcium spike amplitude occurred (Figure 5E). The strongest neuronal responses (i.e. the highest average spike amplitude and frequency) were evoked by magnetic waves at 5, 10 and 20 Hz (Figure 5E and F). Therefore, the 10 Hz frequency was picked as the standard for *in vivo* stimulation in most of the experiments of our study.

**Figure 5.**
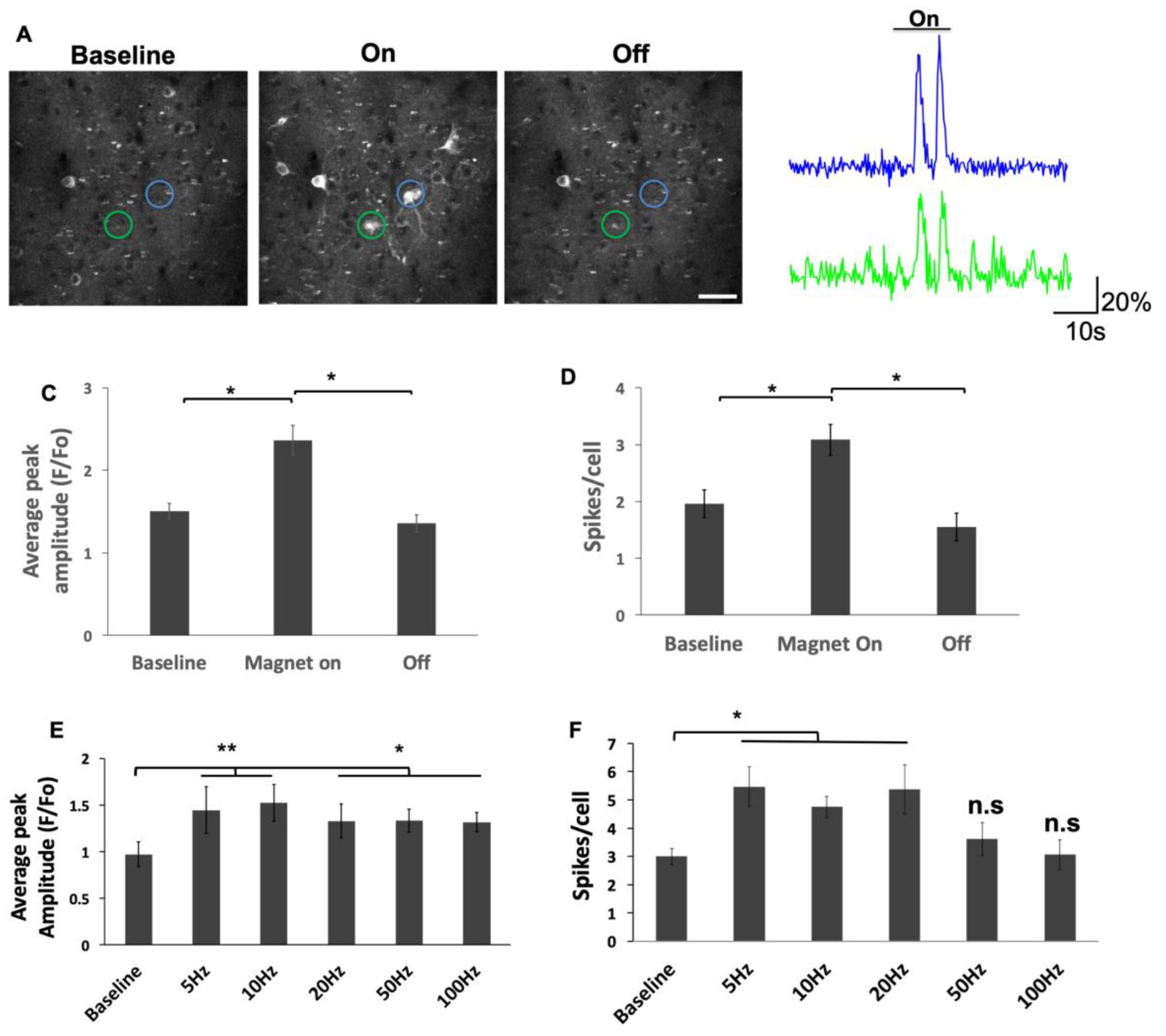
Magnetic stimulation activated cortical layer II/III neuron *in vivo*. **A-B.** Representative images (A) and traces (B) of calcium transients measured from layer II/III cortical neurons. **C-D.** Significant increases in average calcium spike amplitude (C) and mean spike number (D) of cortical neurons after magnet stimulation was turned on. n = 5 mice, *p <0.05, one-way ANOVA, Tukey’s HSD. *<u>Scale bar</u>*: 50 µm. Color of traces correspond to color of analyzed ROI. **E-H.** Increase in average calcium spike amplitude (E, F) and total spikes per cell (G, H) of cortical neurons when magnet was turned on at each frequencies of stimulations. Changes in the amplitudes and spike numbers of calcium transients at electromagnetic stimulation frequencies of 5, 10, 20, 50, and 100 Hz. The data indicate that magnetic waves between 5-10 Hz were most effective in stimulating neurons *in vivo*. n = 5, *p <0.05, **p<0.01, two-way ANOVA, Tukey’s HSD. *<u>Scale bar</u>* – 100um

Multiple brain stimulations over a time period may be required to achieve treatment effect in certain neurological diseases. To evaluate this, we gave animals a single MENs delivery, and then performed MS and calcium imaging at different time points (Figure 6). We found a large increase in the amplitude of calcium spikes when MS is applied immediately after MENs delivery, when compared with baseline where ms was applied without MENs treatment (mean F/F_o_ of 2.7 ± 0.27 vs 1.13 ± 0.07 of baseline, p < 0.05, one-way ANOVA, Tukey’s HSD, Figure 6B, D). Neuronal activity could still be stimulated with similar parameters of MS at 24 hours after MENs delivery (F/F_o_ of 2.35 ± 0.28, p<0.05, one-way ANOVA, Tukey’s HSD), but not at 72 hours after (Figure 6D. F/F_o_ of 1.21 ± 0.18, p>0.05, one-way ANOVA). We also found a higher number of active cells after MENs treatment up to 24 hours (∼90 cells of MENs+MS and 82 cells of 24 hours vs. 62 of ms-only baseline, Figure 6C), and recover to baseline level at 72 hours (∼55 cells, Figure 6C). The result suggests that a single delivery of MENs to brain is effective for non-invasive brain stimulation for at least 24 hours.

**Figure 6.**
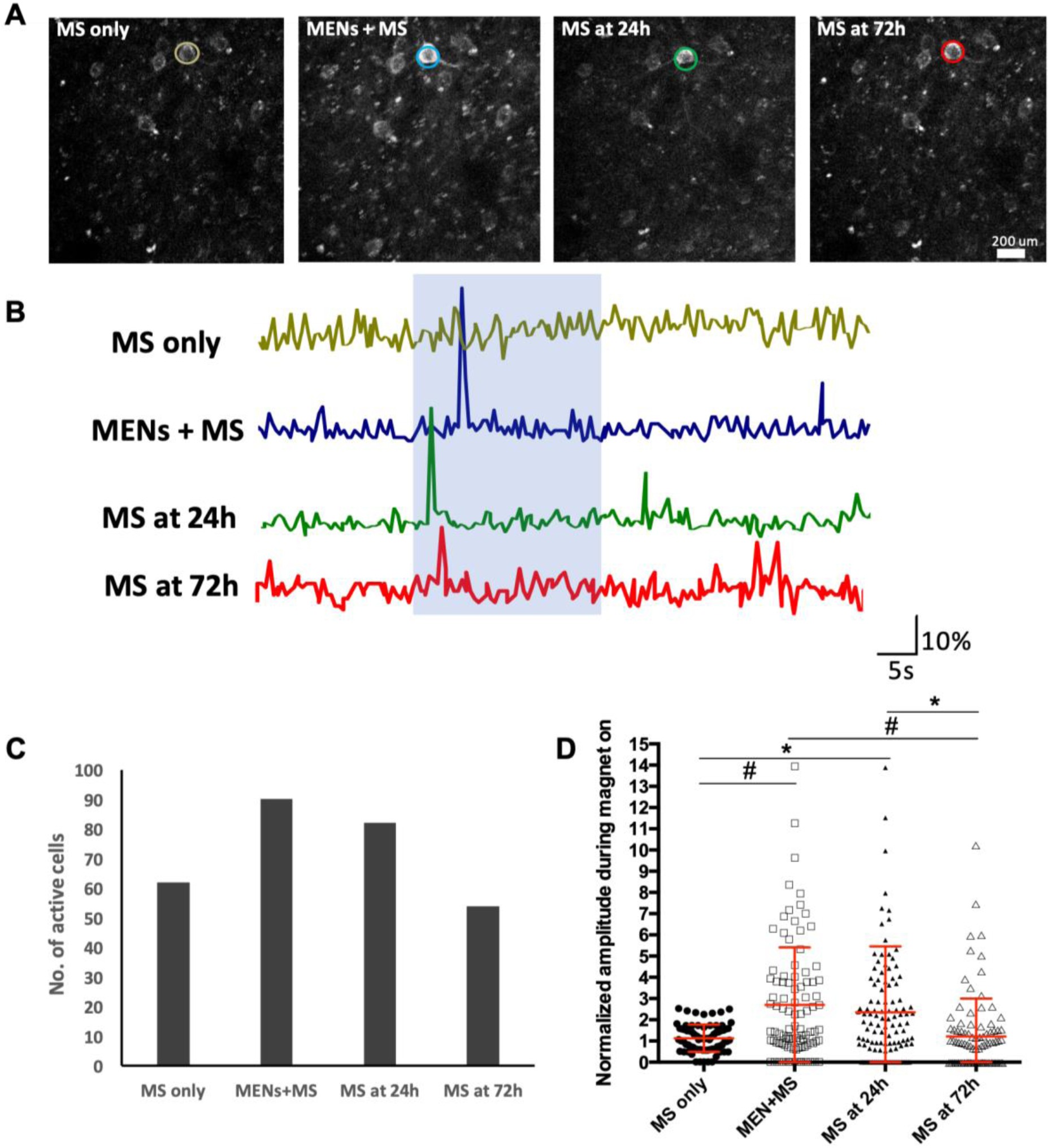
MENs-MS was effective in activating neurons in 24 hours after a single delivery of MENs. **A-B.** Representative two-photon images and traces of calcium signals of same cortical layer II/III neurons at baseline (MS only) and immediately after MENs delivery followed by MS (MEN+MS), MS at 24 hours after, and MS at 72 hours after. 10Hz magnetic stimulation was applied for 20s in the middle of recording period at each time point. **C**. There was an increase in number of active neurons after MENs-MS, which remained higher up to 24 hours after initial delivery. **D.** There were increases in response amplitude immediately after MENs-MS and at least 1 day after initial delivery, which returned to baseline level at 3 days post-MEN delivery, one-way ANOVA, Tukey’s HSD, n=5, *p <0.05, #p <0.005.

To assess the effect of MENs-MS on global network activity, we applied mesoscopic brain imaging technique to evaluate hemispheric calcium response in GCaMP6 transgenic mice (Figure 7A-F). The epicenter region was defined as the location where a permanent magnet was applied to draw MENs to a local cortical region (Figure 7A). Application of a 10 Hz magnetic field induced cortical calcium spikes with significantly higher amplitude and varying latencies in the epicenter and contralateral cortex (p<0.05, repeated ANOVA, Tukey’s HSD, Figure 7 B, C). Comparing the responses among the epicenter and different ipsilateral and contralateral cortical regions revealed a highest peak amplitude at the epicenter and gradual reduction in amplitude (p<0.05, one-way ANOVA, Tukey’s HSD, Figure 7D, E, and F) and an increase in the latency period (Figure 7D, G, and H) at locations further away from the epicenter. The results suggest that neuronal activity was initiated at the epicenter where MENs are localized and spread to more distal brain regions and the contralateral cortex. These data provide strong evidence that MENs-MS can activate and enhance neuronal activity *in vivo*.

**Figure 7.**
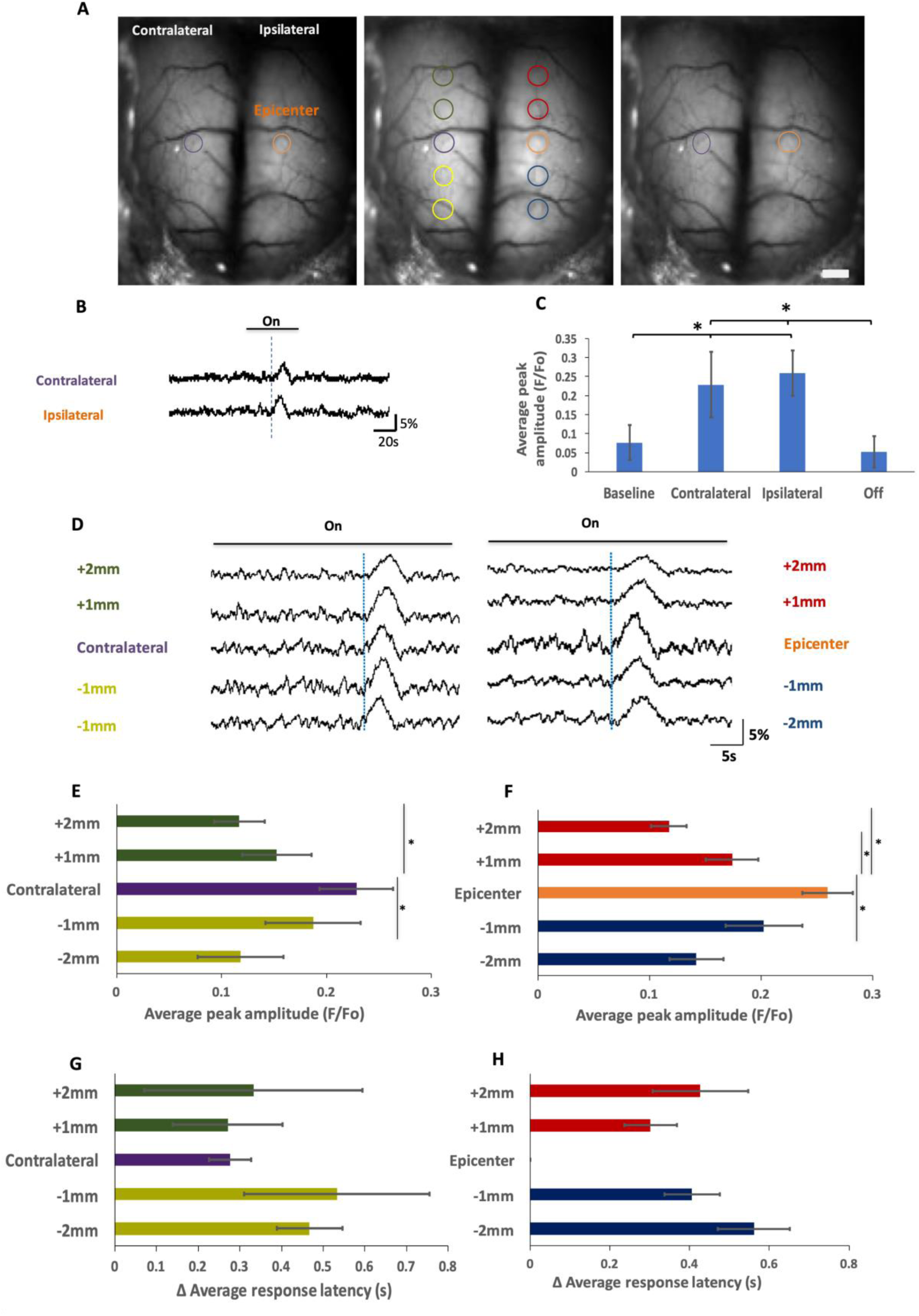
Mesoscopic activity imaging revealed initiation and spreading of MENs-MS induced cortical activity *in vivo*. **A-B.** Representative images (A) and traces (B) of calcium transients measured from both hemispheres at baseline (left image) and after turning on and off magnetic stimulation (middle and right images). The epicenter was a cortical region where a conical magnet was applied to attract MENs. Dotted lines indicate the start of calcium spike. **C.** There were significantly higher amplitudes of calcium signals on cortex ipsilateral (at epicenter region) and contralateral (same region on opposite hemisphere) to MENs delivery induced by MS (10 Hz at 450 Oe for ∼30 seconds). n = 5, *p <0.05, repeated-measured ANOVA, Tukey’s HSD. **D.** Sample traces of calcium transients measured from epicenter, cortical regions anterior and posterior from epicenter, and areas of contralateral cortex. **E, F.** The mean amplitude of calcium signals in the epicenter was higher than that of cortical regions more anterior or posterior to it; the contralateral cortex had similar differences. **G, H.** The latency period at the epicenter was the shortest than all other cortical regions. n = 5, *p <0.05, one-way ANOVA, Tukey’s HSD. *<u>Scale bar:</u>* 1 mm.

### 5. MENs-MS increased cortical c-Fos expression

We found an increase in cortical c-Fos expression only for animals that has both MENs delivery and MS, with an average of 176 ± 27.5 cells/counted region compare to 70.5 ± 15.25 for MENs only group and 51 ± 8 for MS-only group (p<0.05, one-way ANOVA, Tukey’s HSD, Figure 8A, B). MENs only group and MS-only group had similar densities of c-Fos positive cells (Figure 8A, B). The results further support that MENs-MS induced cortical neuronal activity *in vivo*.

**Figure 8.**
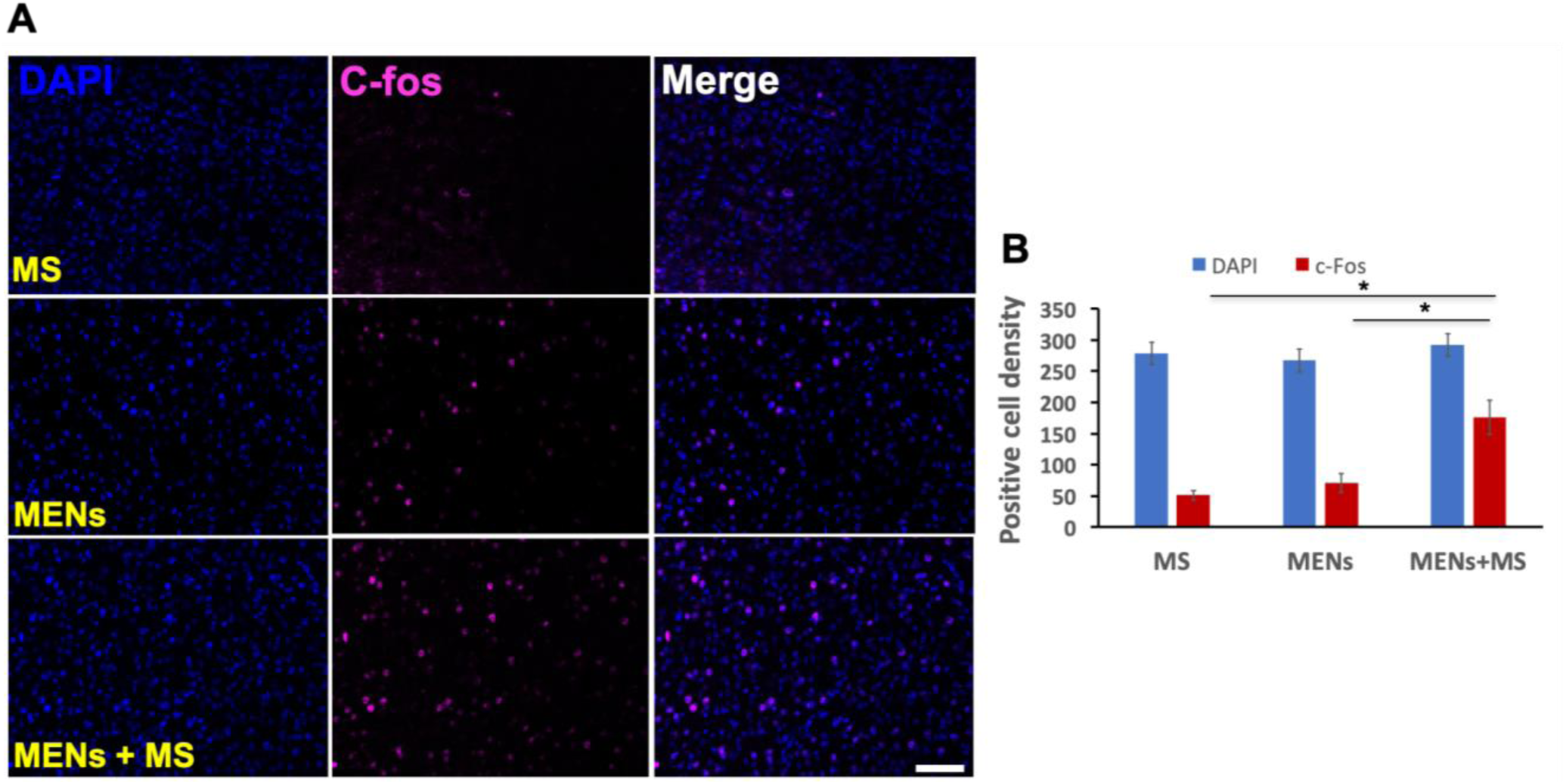
MENs-MS increased the number of c-fos expressing cells. **A.** Representative images of immunofluorescence from mice that received magnetic stimulation (MS, top row), MENs delivery only (middle row), or MENs+MS (bottom row). Images were taken with an 20x objective. **B.** A significantly higher density of c-Fos expressing cells in MENs+MS group than either MS only or MENs only group. *<u>Scale bar:</u>* 20 µm. n=3 mice/group, *: p<0.05, Two-way ANOVA, Tukey’s HSD.

### 6. MENs-MS did not induce astrogliosis and microglial activation

Delivering MENs to a specific brain region may initiate inflammatory responses of glial cells, particularly astrocytes and microglia (Figure 9A-H). To assess whether astrogliosis and microglial activations occurred, we made intravenous injection of 200 µl of MENs at 200 µg/ml and magnetic delivery of MENs. Mice that received MENs delivery and MS were also assessed at various time point after the stimulation. We observed no significant differences in the densities of both IBA1 and GFAP positive cells among all the groups (Figure 9 A-C, G, H). Considering these results together, MENs-MS did not induce any apparent inflammatory response in mice.

**Figure 9.**
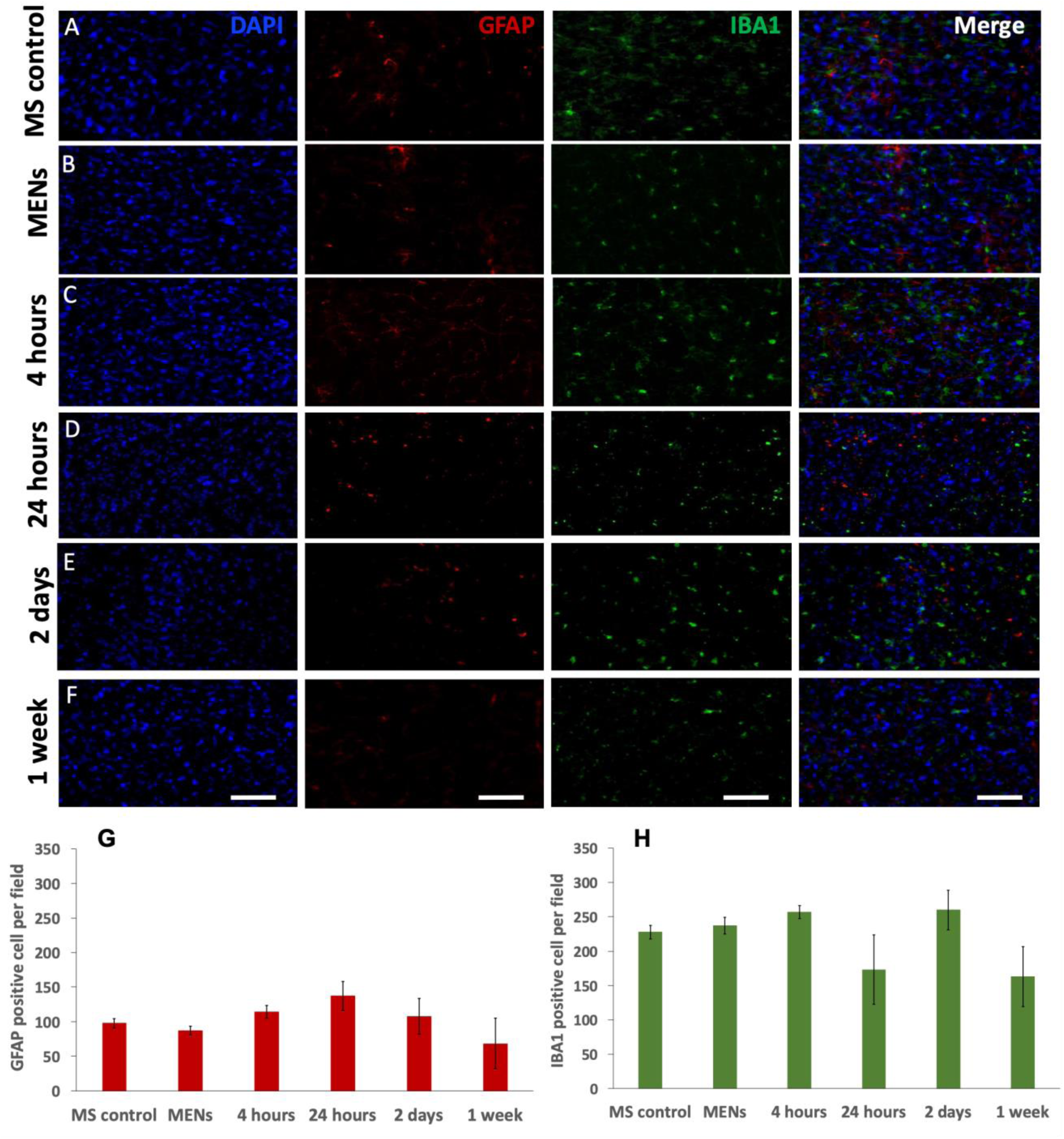
No significant changes in the numbers of GFAP and IBA1 positive cells after MEN delivery and MENs-MS. Brain sections were obtained from six groups of C57BL mice including MS control, MENs-delivery (MENs), and MENs delivery followed by MS at 4h, 24h, 2 days, and 1 week after. **A-F.** Sample confocal images of coronal cortical slices stained for DAPI, GFAP, and IBA1 at different time points. **G, H**. Quantifications of GFAP (G) and IBA1 (H) positive cells showed no significant differences in the numbers of these cells after MEN delivery or followed by MS. n = 4 mice/group. *<u>Scale bar:</u>* 50 µm

## Discussion

In the present study, we investigated the effects of MENs-MS for non-invasive activation of cortical neurons *in vitro* and *in vivo*. By applying an external magnetic field, we non-invasively delivered MENs across the BBB to the brain and wirelessly stimulated neuronal activity with a short latency in a focal cortical region with great efficiency and reliability, without causing detectable neuroinflammatory reaction in the brain.

We developed MENs with a true coreshell nanostructure in which the magnetic core is intrinsically coupled to the piezoelectric shell via perfect lattice matching (Figure 1). To the best of our knowledge, such nanostructures have been fabricated for the first time. Enabling such matching is the key to maximizing the ME coefficient, which in turn is the most important feature of these nanoparticles enabling the creation of an efficient nanotransducer to activate neurons. The small size, on the order of 30 nm, is important not only for their easy penetration through the blood-brain barrier (BBB) but also for ensuring that these nanoparticles can be precisely placed close or directly on the neuronal membrane for maximizing their stimulating capabilities.

Because magnetic stimulation using a pair of electromagnets generates a high level of electromagnetic noise, we used calcium imaging techniques to record neuronal activities instead of using electrophysiological recording. Calcium imaging allowed us to efficiently examine neuronal activities not only in individual neurons *in vivo* and *in vitro* but also in both cortical hemispheres. Upon magnetic application, we observed dramatic increases in neuronal calcium response after a short latency, with about one third of responses having a latency period of <150 ms. Since calcium imaging signals are inherently limited by its slower kinetics than electrical signals and GCaMP6s signal is known to have a temporal resolution of 100-150 ms^21^, the actual latency is estimated to be 50 ms and less. This is a good temporal resolution and a clear indication of the stimulatory effects of MENs on neuronal activity. There were significant increases in both calcium transient amplitude and spike number only during the period when the magnet was turned on. Furthermore, the increases in calcium transient amplitude were seen at all the tested frequencies of magnetic stimulation. Calcium response frequency also elevated significantly, where the most dramatic gains were seen between 5 and 20 Hz of magnet waves.

For in vivo brain stimulation, we first demonstrated that the MENs can be non-invasively delivered to brain parenchyma through intravenous injection followed by application of a permanent magnet on a target region of the skull. The transient existence of fMENs in the blood stream for less than 30 minutes suggests that an effective time window for magnetic delivery of MENs lasted only for less than 30 minutes. We found that the MENs could be drawn across the BBB and localized to a region of the cortical hemisphere where the magnet is applied on the skull. The presence of MENs remained in the brain tissue for at least 24 hours after the delivery. Earlier reports have shown that these MENs, after magnetic delivery to the brain, appear to be localized mainly in the extracellular space both in CSF^26^ and binding to extracellular membranes of various type of CNS residential cells^16, 26^. However, MENs are also found to be distributed in cell cytosol^26^. The chemical structure and integrity of MENs were not compromised during the process of navigation from the blood stream across the endothelial cell layers of the BBB to be localized in the brain^26^. We also assessed the longitudinal profile of MENs effects on neuronal activity and found their stimulatory effects lasted for at least one day after a single treatment and dissipated within three days. This result suggests the majority of the delivered MENs are cleared from the brain within three days, which may reduce potential side-effects or toxicity of the MEN. However, to achieve repetitive brain stimulation over a longer period, additional MEN delivery or modification of the MENs will be required to improve their density or retention. The whole brain mesoscopic imaging revealed activity changes in a spatially dependent manner relative to the location of MENs, suggesting that targeted stimulation of a brain region can be achieved by delivering MENs to a focal brain region defined by the strength of a magnetic field gradient.

One of the major hurdles in nanoparticle-based treatment techniques is the potential toxicity *in vivo* due to nanoparticle agglomerations (e.g. aggregations). Nanoparticle agglomerations could induce severe toxicity and inflammation that may lead to abnormal gene expression, tissue degeneration, and cell death^27, 28, 29^. To prevent particle agglomerations, we coated the surface of MENs with a water-dispersible glycerol mono-oleate (GMO), a well-known water-insoluble and neutral stabilizer that helps prevent aggregates and increases membrane crossing capability^30, 31^. We assessed the neuro-inflammatory response to MENs by evaluating microglial and astrocytic activations at multiple times point after MEN delivery and found no significant changes in microglial activation nor astrogliosis up to one week after MENs delivery and magnetic stimulation. In addition to our findings, Kaushkik et al. also reported no changes in peripheral immune response, liver and kidney toxicity, and motor behavior up to one week after delivering MENs to the cerebral cortex^26^. Furthermore, the MENs have been shown to produce no detectable toxicity on human astrocyte and peripheral blood mononuclear cells at a concentration of 0-200 µg/ml (∼10 ug/kg MENs), which was sufficient to effectively induce an EEG response^16^. Particle functionalization via external coating, such as the SiO_2_ coated - CoFe2O4-PVP (polyvinyl-pyrolidone), was shown to have no toxicity to any vital organs whether it is tissue damage or chromosomal damage^32^. Taken together, these results provide encouraging information on the safety of MENs to the body and brain in rodents and supports its potential application for translational research and clinical treatment.

In addition to imaging neuronal activity, we also stained mice cortical brain slices for c-Fos expression, a classical indicator of activity (e.g. action potential firing)^23^, to confirm the stimulatory effect of MENs on neuronal activity. By comparing sham and MENs-delivery only animals, we found a significant increase in the number of cortical c-Fos expressing cells in MENs-MS animals. Although c-Fos expression in CNS is also observed in some activated astrocytes in inflammatory disease models^33, 34^, our result of no increase in the number of activated astrocytes (Figure 9) suggests that the increased number of c-Fos positive cells are likely activated neurons, which provides histological evidence supporting the effect of MENs-MS on activating neurons. Taken together, these results demonstrate a ground-breaking ability of MENs to enable wireless control of neuronal activity.

Our goal of this study was to demonstrate as a proof-of-concept the ability of MENs to be localized in the brain and efficiently evoke neuronal activity. To further exploit the potential of MENs, additional effort is needed to precisely deliver MENs to a focal region of the brain and to achieve cell-type specific activation. One approach would be to conjugate MENs with an antibody against a cell membrane protein to enable attachment of MENs to a specific cell type or even a subcellular compartment. This approach may allow precise delivery of MENs to excitatory or inhibitory neuronal population. For the past two decades, there have been efforts to enhance delivery of substances, including metal-based magnetic nanoparticles, across the BBB ^35, 36, 37^. One popular way that has been used widely both in research and in clinical application is pre-treatment with BBB-opening compounds such as mannitol^38^ and borneol^39^. Combining these two approaches will help increase MENs delivery to the brain in a more specific manner.

In conclusion, this study demonstrates the ability of using MENs to wirelessly activate individual cortical neurons and cortical network *in vitro* and *in vivo*. We have shown that MENs can be forced to penetrate the BBB to enter brain parenchyma under a magnetic field and we have defined the effective time window and parameters for the delivery. Our *in vitro* and *in vivo* calcium imaging data support that MENs-MS evoke cortical neuronal activity with fast temporal resolution at cellular and global network levels. Furthermore, the process of MENs delivery and magnetic stimulation did not induce detectable neuroinflammation. In contrast to current non-invasive brain stimulation techniques, brain stimulation based on MENs offers a contactless, reliable, and efficient approach to modulate brain activity without the need of genetic modification. With further optimization in MENs delivery and cell- and region-specific targeting, this technique could potentially open a new door to a more robust and precise brain control that currently is not possible.

## Acknowledgements

This publication was made possible with partial support from the N3 program of the DARPA of the Department of Defense (SK, XJ, and LP), the National Science Foundation (NSF) under the grant number ECCS-1935841 (SK and XJ), and from the pre-doctoral fellowship to TN of National Institute of Health grant number NIH-UL1TR002529 (A. Shekhar, PI), National Center for Advancing Translational Sciences, Clinical and Translational Sciences Award and the Indiana University Department of Medicine.

## References

1. Hummel Fea. Effects of non-invasive cortical stimulation on skilled motor function in chronic stroke. Brain 2005, 128: 164–174.

2. Miniussi Cea. Efficacy of repetitive transcranial magnetic stimulation/ transcranial direct current stimulation in cognitive neurorehabilitation.. Brain Stimulation 2008, 1: 326–336.

3. Demirtas-Tatlidede A, …, Pascual-Leone A. Non-invasive brain stimulation in traumatic brain injury. Journal of Head trauma rehabilitation 2012, 27(4): 274–292.

4. Wagner T, Valero-Cabre, A., Pascual-Leone, A. Noninvasive human brain stimulation. Annual Review of Biomedical England 2007, 9: 527–565.

5. Sollmann N, Hauck, T., Tussis, L., Ille, S., Maurer, S., Boeckh-Behrens, T., Ringel, F., Meyer, B., Krieg, S.M. Results on the spatial resolution of repetitive transcranial magnetic stimulation for cortical language mapping during object naming in healthy subjects. BMC Neuroscience 2016, 17(67).

6. Sparing R, Mottaghy, F.M.. Noninvasive brain stimulation with transcranial magnetic or direct current stimulation (TMS/tDCS)-From insights into human memory to therapy of its dysfunction. Methods 2008, 44(f): 329–337.

7. Brunoni AR, Nitsche, M.A., Bolognini, N., Bikson, M., Wagner, T., Merabet, L., Edwards, D.J., Valero-Cabre, A., Rotenberg, A., Pascual-Leone, A., Ferrucci, R., Priori, A., Boggio, P.S., Fregni, F. Clinical research with transcranial direct current stimulation (tDCS): challenges and future directions. Brain Stimulation 2012, 5: 175–195.

8. Gradinaru V. TK, Zhang F., Mogri M., Kay K., Schneider B., Deisseroth K. Targeting and readout strategies for fast optical neural control in vitro and in vivo. Journal of Neuroscience 2007, 27(52): 14231–14238.

9. Fenno L, Yizhar, O., Deisseroth, K.. The development and application of Optogenetics. Annual Review of Neuroscience 2011, 34: 389–412.

10. Chen R, Romero G, Christiansen MG, Mohr A, Anikeeva P. Wireless magnetothermal deep brain stimulation. Science 2015, 347(6229): 1477–1480.

11. Huang H, Delikanli, S., Zeng, H., Ferkey, D.M., Pralle, A. Remote control of ion channels and neurons through magnetic-field heating of nanoparticles. Nature Nanotechnology 2010, 5(6): 602–606.

12. Bauer R, Martin, E., Haegele-Link, S., Kaegi, G., von Specht, M., Werner, B.. Noninvasive functional neurosurgery using transcranial MR imaging-guided focused ultrasound. Parkinsonism & Related Disorders 2014, 20: 175–195.

13. Lipsman N, Schwartz, M. L., Huang, Y., Lee, L., Sankar, T., Chapman, M., et al. MR-guided focused ultrasound thalamotomy for essential tremor: A proof-of-concept study. Lancet Neurology 2013, 12(5): 462–468.

14. Bystritsky A, Korb, A. S., Douglas, P. K., Cohen, M. S., Melega, W. P., Mulgaonkar, A. P., et al. A review of low-intensity focused ultrasound pulsation. Brain Stimulation 2011, 4(3): 125–136.

15. Yue K, Guduru R, Hong J, Liang P, Nair M, Khizroev S. Magneto-electric nano-particles for non-invasive brain stimulation. PloS one 2012, 7(9): e44040.

16. Guduru R, Liang P, Hong J, Rodzinski A, Hadjikhani A, Horstmyer J, et al. Magnetoelectric ‘spin’ on stimulating the brain. Nanomedicine (Lond) 2015, 10(13): 2051–2061.

17. Burgess E, Sylvester M, …, Boggiana MM. Effects of Transcranial Direct Current Stimulations (tDCS) on Binge-eating disorder. International Journal of Eating Disorders 2016, 49(10): 930–936.

18. Guduru R. KS. Magnetic field-controlled release of paclitaxel drug from functionalized magnetoelectric nanoparticles. Particle & Particle Systems Characterization 2014, 31: 605–611.

19. Silasi G. XD, Vanni M., Chen A, Murphy T. Intact skull chronic windows for mesoscopic wide-field imaging in awake mice. Journal of Neuroscience Methods 2016, 267: 141–149.

20. Barson D, Hamodi, A.S., Shen, X. et al. Simultaneous mesoscopic and two-photon imaging of neuronal activity in cortical circuits. Nature Methods 2020, 17: 107–113.

21. Tsai-Wen Chen TJW, Yi Sun, Stefan R. Pulver, Sabine L. Renninger, Amy Baohan, Eric R. Schreiter, Rex A. Kerr, Michael B. Orger, Vivek Jayaraman, Loren L. Looger, Karel Svoboda, Douglas S. Kim. Ultrasensitive fluorescent proteins for imaging neuronal activity. Nature 2013, 499.

22. A. Kerlin MA, V. Berezovskii, R.C. Reid. Broadly tuned response properties of diverse inhibitory neuron subtypes in mouse visual cortex. Neuron 2010, 67(5): 858–871.

23. Bullitt E. Expression of C-fos-Like Protein as a marker for neuronal activity following noxious stimulation in Rats. Journal of Comparative Neurology 1990, 296: 517–530.

24. Imai Y. KS. Intracellular signaling in M-CSF-induced microglia activation: role of Iba1. Glia 2002, 40: 164–174.

25. Liedtke W. EW, Bieri PL., Chiu FC., Cowan NJ., Kucherlapati R., Raine CS. GFAP is necessary for integrity of CNS white matter architecture and long-term maintenance of myelination. Neuron 1996, 17: 607–615.

26. Kaushilk A, …, Nair M. Magnetically guided central nervous system delivery and toxicity evaluation of magneto-electric nanocarriers. Scientific Reports 2016, 6(25309).

27. Lankoff A. SW, Wegierek-Ciuk A., Lisowska H., Refsnes M., Sartowska B., Schwarze P., Meczynska-Wielgosz S., Wojewodzka M, Kruszewski M. The effect of agglomeration state of silver and titanium dioxide nanoparticles on cellular response of HepG2, A549 and THP-1 cells.. Toxicology Letters 2012, 208: 197–213.

28. E F. The role of surface charge in cellular uptate and cytotoxicity of medical nanoparticles. International Journal of Nanomedicine 2012, 7: 5577–5591.

29. Okuda-Shimazaki J. TS, Kanehira K., Sonezaki S., Taniguchi A. Effects of Titanium Dioxide nanoparticle aggregate size on gene expression. International Journal of Molecular Science 2010, 11: 2383–2392.

30. Nair M, Guduru R., Liang P., Hong J., Sagar V., Khizroev S. Externally ocntrolled on-demand release of anti-HIV drug using magneto-electric nanoparticles as carriers. Nature Communications 2013, 4(1707).

31. Dilnawaz F. SA, Mewar S., Sharma U., Jagannathan N.R., Sahoo S. The transport of non-surfactant based paclitaxel loaded magnetic nanoparticles across the blood brain barrier in a rat model. Biomaterials 2012, 33(33): 2936–2951.

32. Kim JS, Yoon T, …, Cho MH. Toxicity and tissue distribution of magnetic nanoparticles in mice. Toxicological Sciences 2006, 89(1): 338–347.

33. Wu B. YA. Quercetin inhibits c-fos, Heat Shock Protein, and Glial fibrillary acidic protein expression in injured astrocytes. Journal of Neuroscience Research 2000, 62: 730–736.

34. Groves A. KY, Jonnalagadda D., Rivera R., Kennedy G., Mayford M., Chun J. A functionally defined in vivo astrocyte population identified by c-Fos activation in a mouse model of multiple sclerosis modulated by S1P signaling: immediate-early astrocytes (ieAstrocytes). eNeuro 2018.

35. Koziara JM LP, Allen DD, Mumper RJ. The blood-brain barrier and brain drug delivery. Journal of Nanoscience Nanotechnology 2006, 6: 2712–2735.

36. Y. J. Yu YZ, M. Kenrick, K. Hoyte, W. Luk, Y. Lu, J. Atwal, J. M. Elliott, S. Prabhu, R. J. Watts, M. S. Dennis.. Boosting brain uptake of a therapeutic antibody by reducing its affinity for a transcytosis target. Science Translational Medicine 2011, 3.

37. Qiao R. JQ, Huwel S., Xia R., Liu T., Gao F., Galla H., Gao M. Receptor-mediated delivery of magnetic nanoparticles across the blood-brain barrier. ACS Nano 2012, 6(4): 3304–3310.

38. Chu C. JA, Lesniak W., Thomas A., Lan X., Linville R. et al. Optimization of osmotic blood-brain barrier opening to enable intravital microscopy studies on drug delivery in mouse cortex. Journal of Controlled Release 2020, 317: 312–321.

39. Zhang Q. FB, Zhang Z.. Borneol, a novel agent that improves central nervous system drug delivery by enhancing blood-brain barrier permeability. Drug Delivery 2017, 24(1): 1037–1044.

